# A molecular inversion probe and sequencing-based microsatellite instability assay for high throughput cancer diagnostics and Lynch syndrome screening

**DOI:** 10.1101/382754

**Authors:** Richard Gallon, Christine Hayes, Lisa Redford, Ghanim Alhilal, Ottie O’Brien, Amanda Waltham, Stephanie Needham, Mark Arends, Anca Oniscu, Angel Miguel Alonso, Sira Moreno Laguna, Harsh Sheth, Mauro Santibanez-Koref, Michael S Jackson, John Burn

## Abstract

**Background:** Clinical guidelines recommend microsatellite instability (MSI) and *BRAF* V600E testing of all colorectal cancers (CRCs) to screen for Lynch syndrome (LS), a hereditary predisposition to cancer. MSI is also associated with response to immunotherapy. However, uptake of MSI testing is poor and current assays are not suitable for high throughput diagnostics.

We aimed to develop a cheap and scalable sequencing assay for MSI classification, which is robust to variables in clinical samples and simultaneously tests for *BRAF* V600E to streamline the LS screening pipeline.

**Methods:** 24 short (7-12bp) microsatellites and the *BRAF* V600E locus were amplified in multiplex using single molecule molecular inversion probes (smMIPs) and sequenced using the Illumina MiSeq platform. Reads were aligned to reference genome hg19. An MSI classifier was trained from 98 CRCs and validated in 99 independent CRCs collected in pathology laboratories in Edinburgh, Spain and Newcastle.

**Results:** The smMIP-based MSI assay has 100% accuracy for MSI status relative to MSI Analysis System (Promega). MSI classification is reproducible (100% concordance) and is robust to sample variables, detecting less than 5% MSI-high content in template DNA and giving reliable classification from sequencing only 75 DNA molecules per marker. *BRAF* V600E was detected with mutant allele frequencies down to 1.7%.

**Conclusions:** Our short microsatellite, smMIP-based, MSI assay provides a cheap and fully automatable assessment of MSI status and *BRAF* mutation. It is readily scalable to high throughput cancer diagnostics, and is suitable both as a companion diagnostic for immunotherapy and for streamlined LS screening.

## Introduction

Colorectal cancer (CRC) is the third most common cancer in Western society^1^.

Approximately 1 in 6 CRCs have genome-wide insertion-deletion mutation in >30% of microsatellites, defined as high microsatellite instability (MSI-high)^2^, a phenotypic manifestation of deficiency of the mismatch repair (MMR) system^3^. Lynch syndrome (LS) is a cancer-predisposition syndrome caused by germline pathogenic variants affecting one of four MMR genes, with a lifetime cancer risk up to 70%^4^. The LS cancer spectrum includes CRC, endometrial cancer and others. Clinical management of LS includes 1-2 yearly surveillance colonoscopy, prophylactic surgery^5^, and chemoprevention using daily aspirin^6^. Following the two hit hypothesis of tumour progression, nearly all LS CRCs are MMR deficient (MMRd)^7^. Guidelines from the UK National Institute of Health and Care Excellence^8^ and others^9^ advocate that all CRCs should be tested for MMR deficiency to screen for LS. *BRAF* V600E or *MLH1* methylation testing increases the specificity of LS screening by removing sporadic MMRd CRCs^10,11^. MMR deficiency can also inform choice of therapy as affected CRCs are sensitive to immune checkpoint blockade^12^, which is FDA-approved as a second line therapeutic for any MSI-high solid cancer^13^.

The dominant MMR deficiency tests are MSI detection by PCR fragment length analysis (FLA), or assessment of MMR protein expression by immunohistochemistry (IHC). FLA using panels exclusively composed of mononucleotide repeats (MNRs) achieves sensitivity and specificity of 97% and 100% respectively^14^ and IHC of all 4 MMR proteins has 93% sensitivity and 95% specificity^15^. IHC misses approximately 5% of MMRd CRCs^14^, due to mutations that lead to loss of MMR function but retain antigenicity^15^. A high accuracy, diagnostic test should also be robust to sample variables, be reproducible and meet clinical demands such as rapid turnaround time and low cost. FLA has 98% reproducibility between independent laboratories^16^, whilst IHC shows greater heterogeneity in results interpretation^15^. Both are considered accurate for tumour cell content >10%^17^ and the National Institute of Health Research Health Technology Assessment found both to be cost-effective within the LS screening pipeline, with incremental cost-effectiveness ratios <£20,000 per quality-adjusted life year gained^18^. However, testing all CRCs to screen for LS, and predictive testing of any cancer for response to immune checkpoint blockade therapy, will require scalable assays. IHC and FLA are limited by reliance on case-by-case result analysis, with IHC needing trained pathologists to assess variable staining patterns^15^ and FLA needing experienced operators to interpret PCR “stutter” peaks generated by the error-prone markers used^16^. Due to the unsuitability of current assays for high throughput, the uptake of MMR deficiency testing has been poor: only 28% of 152,993 CRC cases were tested between 2010-2012 in the USA^19^ despite concurrent estimation that only 1.2% of the LS gene carrier population was known^20^ and over a decade of testing recommendations^9^.

Next generation sequencing (NGS) of tumours to diagnose MSI have been developed with sensitivities and specificities of >95%^21^. Tumour-sequencing could reduce the recommended LS screening pipeline to two steps (tumour-sequencing and germline confirmation) by simultaneous detection of MSI, *BRAF* V600E and MMR gene mutations^22^. Coupled tumour and germline-sequencing can also determine the somatic origin of MMR deficiency in 52% of Lynch-like tumours with double somatic MMR mutations, which may avoid unnecessary management of these CRCs as LS^23^. However the cost of tumour-sequencing is a barrier to its deployment, with an estimate of 607±207€ per sample in a recent French, nationwide study of tumour-sequencing in clinical practice ^24^. Cheaper assays that target multiple, clinically actionable markers are needed. MSIplus, for example, is a targeted NGS-based assay analysing MSI, *BRAF* and *RAS* gene mutations, but includes long (up to 28bp) and error prone microsatellite markers^25^. Furthermore, the robustness of NGS-based MSI assays to multiple sample variables are rarely or inadequately assessed, with a lack of quality control to ensure results reliability, which is desirable for deployment of assays into the clinic^26^.

We have recently published a PCR and NGS-based assay for MSI that utilises a novel set of short (7-12bp) and monomorphic MNRs, selected to reduce PCR and sequencing error and ease interpretation relative to longer markers^27,28^. SNPs neighbouring each of the microsatellites are used to determine allelic bias of instability in heterozygote patients, increasing signal-noise discrimination. A naïve Bayesian MSI classifier was developed for these markers which has ≥97% sensitivity and specificity relative to microsatellite FLA^29^. Here, we create a fully automatable, modular, and cheap MSI assay, suitable for high throughput MMR diagnostics and two-step screening for LS, by multiplex amplification of our markers and the *BRAF* V600E locus using single molecule molecular inversion probe (smMIP) technology^30^. smMIPs have previously been used to multiplex large panels of long (16-40bp) microsatellites^31^, and read-tagging with molecular barcodes provides a count of template molecules sequenced as a quality control^32^. We show that our smMIP-based MSI assay has 100% sensitivity and specificity for MMR deficiency relative to FLA, and detects *BRAF* V600E mutations missed by conventional techniques. MSI calling is reproducible and robust to sample variables, with accurate classification of DNA equivalent to 3% MMRd tumour cell content and from a minimum sequencing depth of 75 template DNA molecules per marker.

## Materials and Methods

### Samples

19 CRC DNAs were provided by the Department of Molecular Pathology, University of Edinburgh, UK. 73 CRC DNAs were provided by the Genetics Service of the Complejo Hospitalario de Navarra and Hereditary Cancer Group, IDISNA, Biomedical Research Institute of Navarra, Spain. These 92 samples were originally from FFPE tissue and were residual stocks remaining from Redford *et aß^9^.* These were used in the MSI classifier training cohort.

105 CRC DNAs or CRC FFPE tissue samples were provided by the Northern Genetics Service, Newcastle Hospitals NHS Foundation Trust, UK, after ethical review (REC reference: 13/LO/1514). 6 samples were included in the MSI classifier training cohort and the remaining 99 were used for classifier validation. 46 of the MMRd samples were tested for *BRAF* V600E by high resolution melt curve (HRM) analysis^33^ on a LightCycler 480 (Roche).

19 DNAs extracted from peripheral blood lymphocytes from patients with no evidence of familial cancer syndromes, consented for sample use for assay development, were used as microsatellite stable (MSS) controls.

All CRC samples were independently tested for MMR deficiency by MSI Analysis System v1.2 (Promega) in the contributing pathology laboratory.

All samples were anonymised by the contributing laboratories.

### Cell Lines and Cell Culture

DNA from embryonic stem cell H9 was a gift from L. Lako (Newcastle University, UK), and was used as an MSS control.

Both HCT116 and K562 cells were gifted by the Irving Research Group (Newcastle University, UK). HCT116 CRC cell line, containing hemizygous *MLH1* truncation, was used as an MSI-high control. K562 chronic myeloid leukaemia cell line was used as an MSS control. HCT116 and K562 cells were grown in RPMI growth medium containing 2mM L-glutamine (Gibco), 10% fetal bovine serum (Gibco), 60μg/ml penicillin and 100μg/ml streptomycin (Gibco) at 37°C and 5% CO2. HCT116 cells were split and harvested at 80-90% confluence by decanting expired growth medium, washing in 5ml PBS (Gibco) and detaching cells using 0.05% Trypsin-EDTA (Gibco). K562 cells were split and harvested at a density of 1×10^6^cells/ml.

### DNA Extraction and Quantification

DNA was extracted from FFPE CRC tissue using the GeneRead DNA FFPE Kit (QIAGEN).

DNA was extracted from cell lines using the Wizard Genomic DNA Purification Kit (Promega).

DNAs were quantified using QuBit 2.0 Fluorometer (Invitrogen) and QuBit dsDNA BR/HS Kits (Invitrogen).

### Marker Selection

A total of 24 MNRs with length 7-12bp and neighbouring SNP^29^ were selected to test for MMR deficiency. *BRAF* V600E was included to screen for sporadic MMRd CRCs.

### smMIP Design

MIPgen^34^ was used to generate smMIPs for each marker using reference genome hg19, indexed with SAMtools v1.3 and BWA v0.7.12, and a bed file of marker loci (Supplement S1). MIPgen parameters were: tag size 6,0, minimum capture size 120, and maximum capture size 150.

Final smMIP designs (Supplement S2) were selected by the following criteria: successful capture of marker and associated SNP (where applicable), no SNPs in the smMIP extension or ligation arms, logistic score >0.8, and successful generation of smMIP amplicons of the expected size.

All smMIPs, smMIP amplification primers, and custom sequencing primers (Supplement S2) were synthesised by and purchased from Metabion.

### smMIP Phosphorylation

smMIPs were individually phosphorylated using 10U of T4 Polynucleotide Kinase (NEB), 1X T4 DNA Ligase buffer (NEB) and 1μM of unphosphorylated smMIP in a 100μl reaction volume, and incubated at 37°C for 45 minutes and 80°C for 20 minutes. Phosphorylated smMIPs were diluted 1:10,000 using TE buffer (Sigma) in a multiplex pool, such that each smMIP was at 0.1nM (0.1fmol/μl).

### smMIP Amplification

smMIP-multiplexed amplification was based on Hiatt *et al^30^* using a SensoQuest thermocycler (SensoQuest GmbH), with minor modifications to the protocol: Herculase II Polymerase (Agilent) was used during gap-fill and amplification steps, and amplification thermocycling used 98°C for 2 minutes, 30 cycles of 98°C for 15 seconds, 60°C for 30 seconds, and 72°C for 30 seconds, followed by 72°C for 2 minutes. smMIP reaction products (240-270bp) were analysed using agarose gel electrophoresis (3% gel run at 80mV for 60 minutes) or QIAxcel (QIAGEN) using method AL420.

### Library Preparation and Sequencing

Sequencing libraries were prepared by purification of smMIP amplicons using Agencourt AMPure XP Beads (Beckman Coulter), diluting purified amplicons to 4nM in 10mM Tris pH 8.5 and pooling in equal volumes.

Libraries were sequenced using the MiSeq platform (Illumina) with the GenerateFastq workflow, paired end sequencing, and smMIP custom sequencing primers^30^.

### Sequencing Read Analysis and MSI Classification

Sequencing reads from MiSeq-generated fastq files were aligned to hg19 using BWA v0.6.2^35^. Variants in microsatellite length, SNP and mutation hotspot loci were identified from .sam files, and only variants observed on both reads of a pair were tabulated by custom R scripts.

The MSI classifier used custom R scripts and the algorithm previously described by Redford *et al^29^.* In summary, to determine the relative probability that a sample is modelled by an MMRd versus an MMRp phenotype, both the proportion of reads containing microsatellite deletions and the allelic bias of deletions are used. Scores >0 classify a sample as MMRd (MSI-high). The algorithm is trained in one cohort of samples and validated in a second, independent cohort.

Custom R scripts were used for all other analyses of reads and variants (available upon request).

### Statistics and Graphics

Statistical tests and graph generation was performed in R (v3.3.1), utilising R package ggplot2.

## Results

### MSI Status is Accurately and Reproducibly Classified

To train the MSI classifier algorithm, 24 microsatellite and *BRAF* V600E markers were amplified from 51 MMRd CRCs, 47 MMRp CRCs, and HCT116 and H9 cell lines as MSI-high and MSS controls (see methods). Amplicons were sequenced to a mean read depth (±sd) of 3719±3149 reads/marker/sample with 75.3% of base-calls ≥Q30. The trained MSI classifier subsequently typed these same samples with 100% sensitivity (95% CIs: 93.0-100.0%) and 100% specificity (95% CIs: 92.5-100.0%) (Figure 1A). Classification by the most discriminatory 6 microsatellites also achieved 100% accuracy, suggesting marker redundancy (Figure 1A).

**Figure 1:**
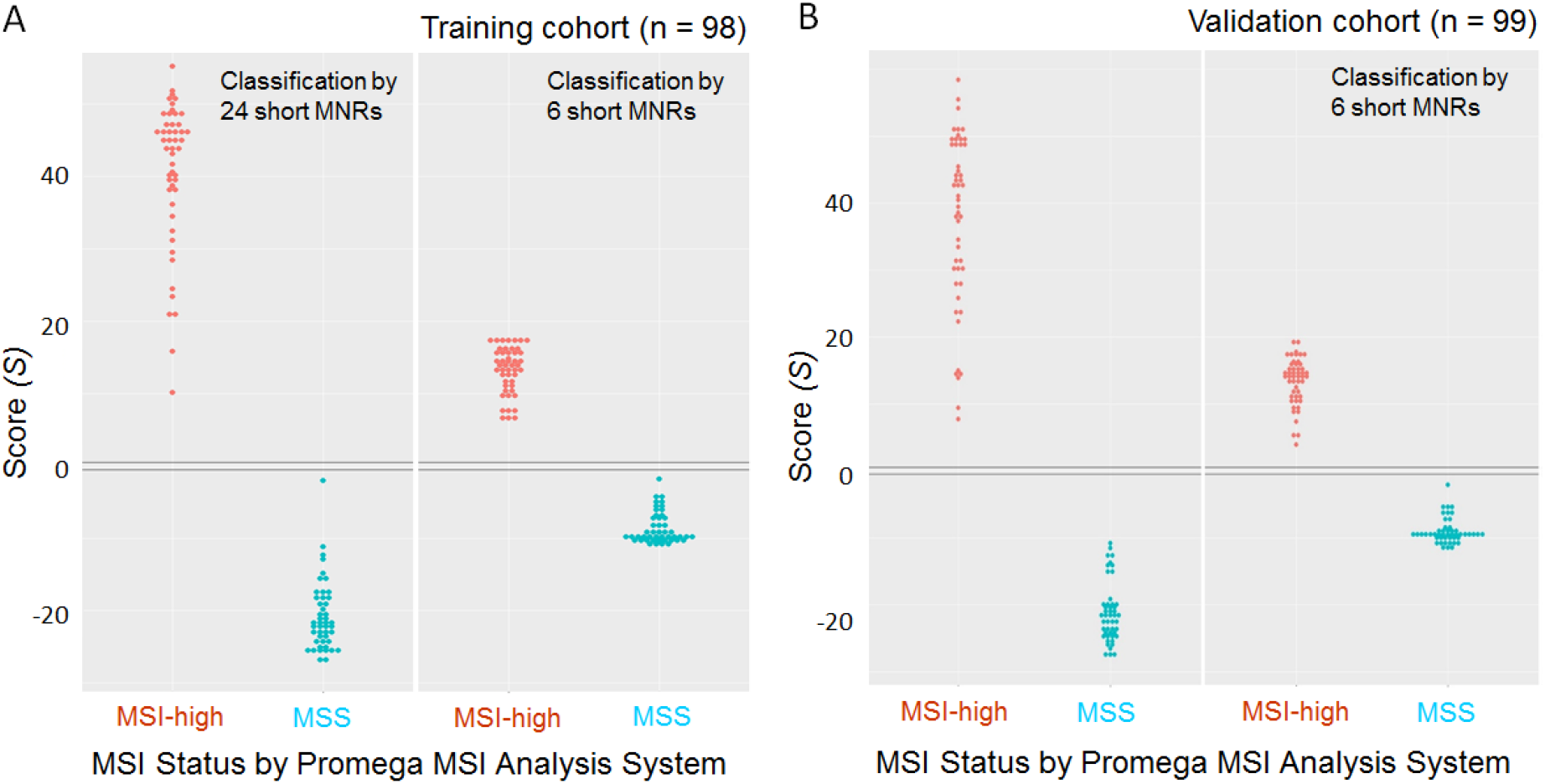
MSI classification of CRCs using a smMIP-based sequencing assay. The MSI classifier is able to type CRCs with 100% sensitivity and 100% specificity using a panel of 24 microsatellites (left hand panels) or only 6 microsatellites (right hand panels), relative to independent assessment by MSI Analysis System vl.2 (Promega), in both **(A)** the training cohort (n = 98) and **(B)** the validation cohort (n = 99). Scores > 0 are classified as MSI-high, Scores < 0 are classified as MSS.

A read-balanced smMIP pool, based on per marker read depths from the training cohort, was created to equalise the number of reads generated from each marker (Supplement S3). 50 MMRd CRCs and 49 MMRp CRCs were then amplified as an independent validation cohort (see methods), and sequenced to a mean read depth (±sd) of 7320±4192 reads/marker/sample with 57.2% of base-calls ≥Q30. The MSI classifier achieved 100% sensitivity (95% CIs: 92.9-100.0%) and 100% specificity (95% CIs: 92.8-100.0%) using either 24 microsatellites or the 6 most discriminatory microsatellites identified from the training cohort (Figure 1B).

To assess assay reproducibility, 16 MMRd and 16 MMRp CRCs from the validation cohort were amplified a second time using a new read-balanced smMIP pool, again targeting the 24 microsatellite markers and *BRAF* V600E. These amplicons were sequenced to a mean read depth (±sd) of 5408±2160 reads/marker/sample with 85.4% of base-calls ≥Q30. Classification was 100% concordant with previous results and classifier scores were strongly correlated between sample repeats (β = 0.97, p < 10^-16^, R^2^ = 0.97).

### *BRAF* V600E is Detectable at Low Variant Frequencies

All of the 14 CRCs that tested positive for *BRAF* V600E by HRM had ≥5% mutant reads assigned to the *BRAF* V600E locus using the smMIP-based sequencing assay. Of the 32 CRCs that tested negative for *BRAF* V600E by HRM, 30 samples had *BRAF* V600E detected in ≤0.6% of reads and 2 samples had *BRAF* V600E detected in 1.67% and 1.72% of reads, suggesting these 2 samples may contain true mutations not detected by HRM.

Using a ≥1% mutant read threshold, *BRAF* V600E was detected in 9.4% of MMRp CRCs (95% CI: 4.4-17.1%) and in 36.6% of MMRd CRCs (95% CI: 27.3-46.8%). 100% concordance was observed between BRAF V600E mutation calling in the repeat testing of 16 MMRd and 16 MMRp CRCs, with strong correlation of the proportion of mutant reads detected (β = 0.93, p < 10^-16^, R^2^ = 0.99).

### MSI Classification is Robust to Low MSI-high Content

To assess the lower limit of detection (LLoD), defined here as the lowest proportion of MSI-high DNA within total template DNA at which a sample is classified as MSI-high, a DNA-mixture series of 0.78-100% MSI-high DNA content (log2 increments) was created in triplicate, by mixing HCT116 MSI-high DNA into control MSS DNA extracted from peripheral blood leukocytes. This triplicate series and control MSS DNAs were amplified using a read-balanced smMIP pool. Amplicons were sequenced to a mean read depth (±sd) of 4763±1288 reads/marker/sample with 84.7% of base-calls ≥Q30.

Increasing the MSI-high DNA content of the template DNA increased the proportion of reads containing insertion-deletion mutations in the microsatellite (Figure 2A); the observed and the expected proportions (Supplement S4) were strongly correlated (β = 1.009, p = 2×10^-16^, R^2^ = 0.996, Figure 2B), giving confidence in the accuracy of the DNA-mixture series. MSI classification of the DNA-mixture series was accurate from 3.13% or more MSI-high content in each replicate series (Figure 2C), approximating the LLoD to 3%. To compare with FLA, replicates ranging from 1.56-12.5% MSI-high DNA content were independently classified using the MSI Analysis System (Promega), with the observer blinded to both sample content and experimental purpose. FLA reliably detected 6.25% MSI-high DNA content (Table 1).

**Table 1:** Microsatellite instability classification by fragment length analysis of DNA-mixtures of varying MSI-high DNA content. A series of samples with varying MSI-high DNA content were analysed using the MSI Analysis System (Promega). Fragment length analysis correctly classified samples as MSI-high when they contained ≥3.13% MSI-high DNA. However, at 3.13% MSI-high DNA content the pathologist was uncertain of the status of all 5 markers. Therefore, confident classification as MSI-high was only achieved in samples with ≥6.25% MSI-high DNA content. These same samples were analysed using our smMIP and NGS-based MSI assay.

| MSI-high content (%) | Diagnosis | Unstable Markers | Uncertain Markers |
| --- | --- | --- | --- |
| 1.56 | MSS | 0/5 | 0/5 |
| 1.56 | MSS | 0/5 | 0/5 |
| 1.56 | MSS | 0/5 | 0/5 |
| 3.13 | MSI-high | 3/5 | 5/5 |
| 3.13 | MSI-high | 2/5 | 5/5 |
| 3.13 | MSI-high | 2/5 | 5/5 |
| 6.25 | MSI-high | 5/5 | 2/5 |
| 6.25 | MSI-high | 5/5 | 0/5 |
| 6.25 | MSI-high | 5/5 | 0/5 |
| 12.5 | MSI-high | 5/5 | 0/5 |
| 12.5 | MSI-high | 5/5 | 0/5 |
| 12.5 | MSI-high | 5/5 | 0/5 |

**Figure 2:**
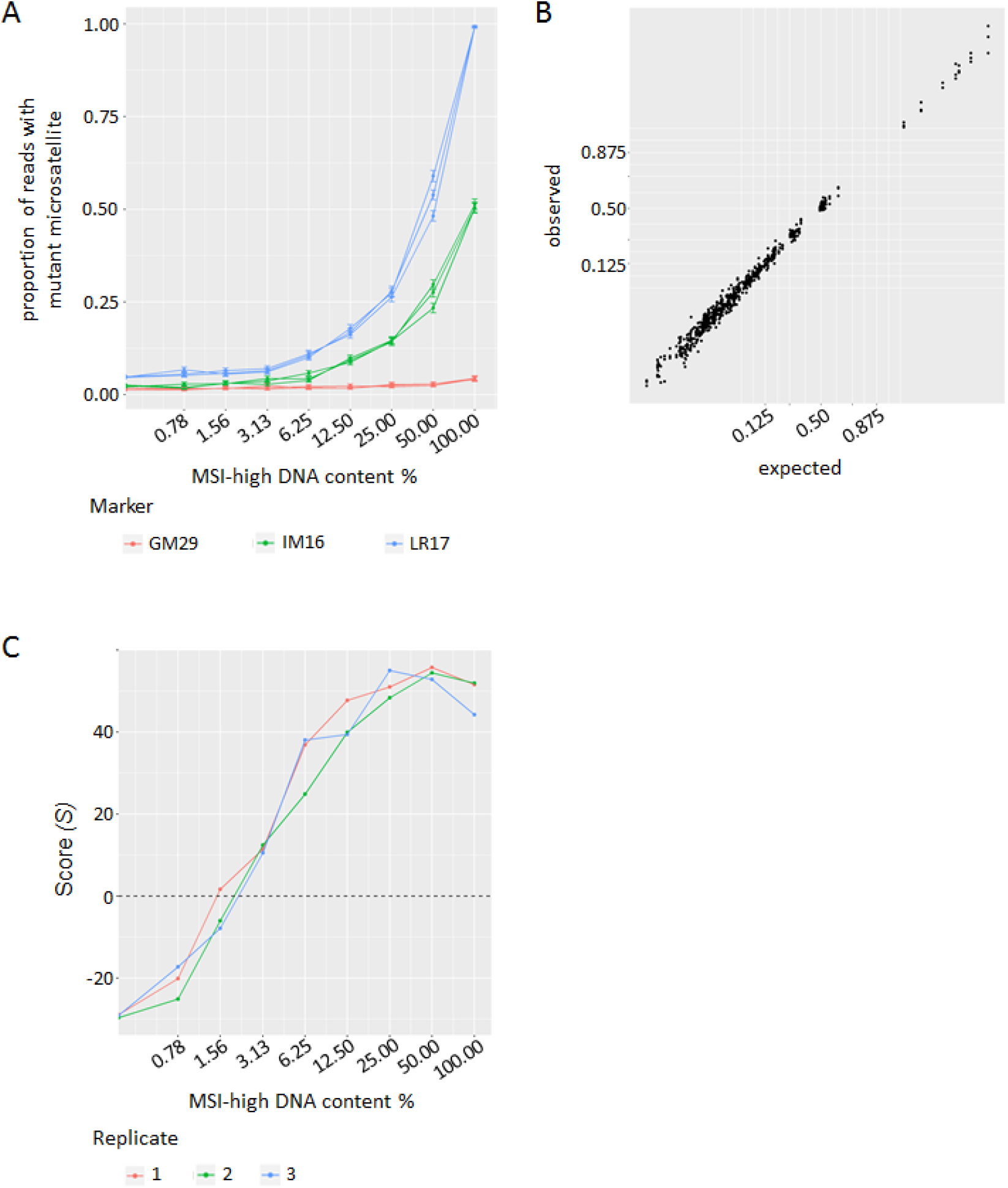
Assessing the lower limit of detection (LLoD) of the assay. **(A)** The proportion of reads containing insertion-deletion mutations in microsatellites increases as the MSI-high DNA content increases in the sample. This is shown for three markers: GM29, representative of markers that are un-mutated or sub-clonally mutated in the MSI-high DNA; IM16, representative of markers that are heterozygous mutated in the MSI-high DNA; and LR17, representative of markers that are homozygous mutated in the MSI-high DNA. **(B)** Across DNA-mixture samples of varying MSI-high DNA content, the observed and the expected proportions of reads with a mutant microsatellite correlate very closely with expected values. **(C)** The MSI classifier calls MSI-high with 100% accuracy in samples with ≥3.13% MSI-high DNA content.

### MSI Classification is Reliable from sequencing 75 Molecules per Marker

To establish the lowest quantity of template DNA required for accurate smMIP-based classification, 2-fold dilution (0.78-100ng) series of 9 DNA samples, comprising 3 cell lines (HCT116, K562 and H9), 3 MMRd CRCs and 3 MMRp CRCs, were amplified using a read-balanced smMIP pool. Samples were selected based on availability of residual DNA. Amplicons generated from 3.13-100ng of template DNA, selected by visual detection of the reaction products (Supplement S5), were sequenced to a mean read depth (±sd) of 243,073±64,485 reads/sample with 82.8% of base-calls ≥Q30.

The number of template molecules sequenced, as measured by the number of molecular barcodes detected, was compared to the input quantity of DNA of the 9 samples, and the two were closely correlated (β = 0.84-0.96, p < 10^-3^, R^2^ = 0.986-0.997, Figure 3A), giving confidence in the dilution series. The MSI classifier accurately typed samples using ≥12.5ng of template DNA (Figure 3B). However, two MMRd CRC samples (207950 and 244881) showed large changes in classifier score between 12.5ng and 25ng of template DNA. These samples were derived from FFPE tissue, suggesting that low quality DNA may be responsible for the variation in classifier score. This was supported by the lower number of molecular barcodes detected in these two samples (Figure 3A). Comparison of classifier scores with the mean number of molecular barcodes detected suggested that sequencing a minimum of 75 template molecules per marker is sufficient to reliably classify these samples (Figure 3C).

**Figure 3:**
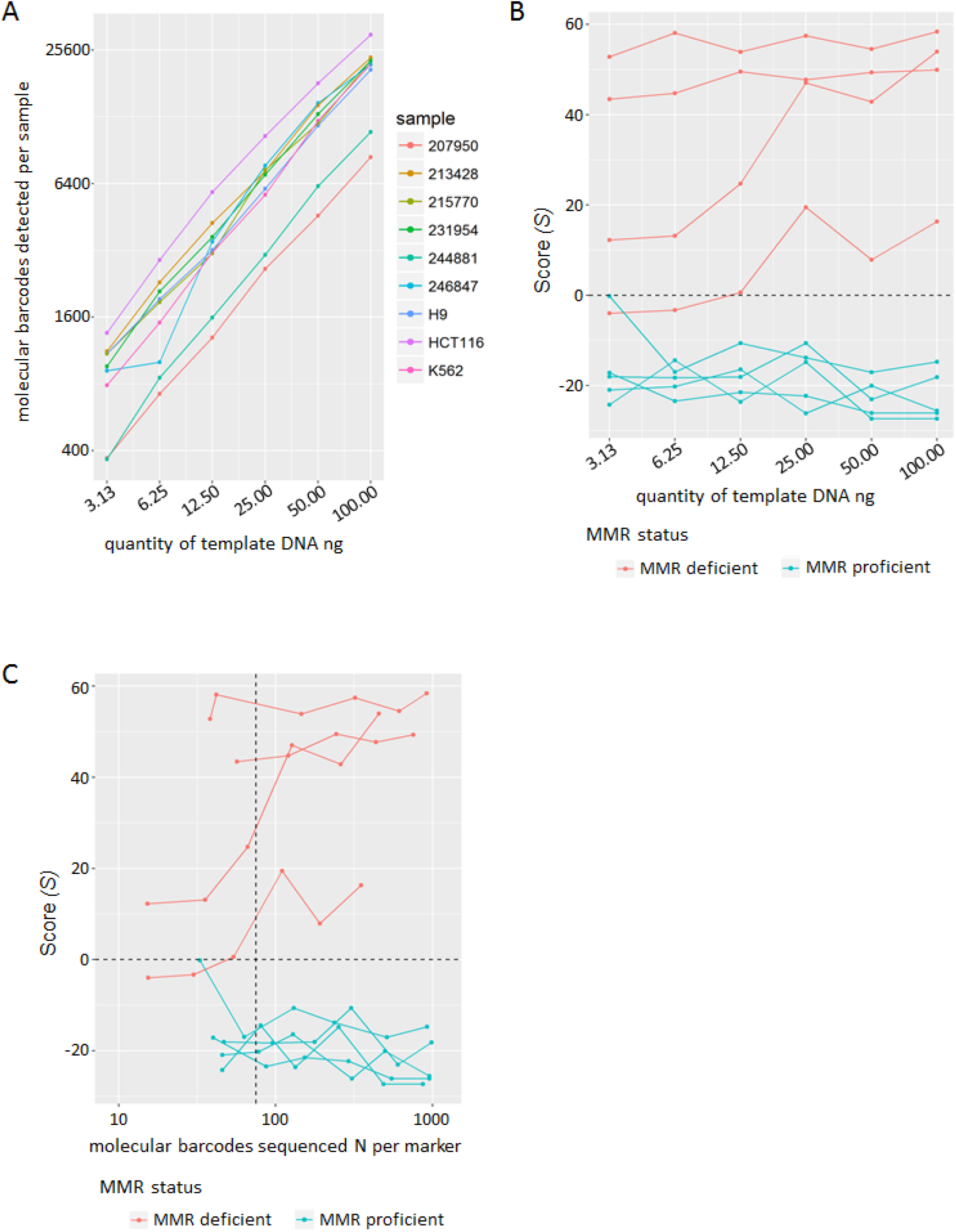
Assessing assay robustness to low quantity and quality of DNA sample. **(A)** The sum of molecular barcodes detected per sample represents the number of template DNA molecules successfully sequenced across all of the markers. The number of molecular barcodes correlates with the amount of template DNA used for each sample. **(B)** The MSI classifier correctly classifies all 9 samples using ≥12.5ng of template DNA. **(C)** A minimum of 75 molecular barcodes per marker (vertical dotted line) ensures correct sample classification.

## Discussion

Our smMIP-based MSI assay uses monomorphic and short (7-12bp) MNRs that have significantly lower PCR and sequencing error rates compared to longer markers^27^, including those used by the MSI Analysis System (Promega) and NGS-based assays^25,31^. Automated MSI classification from sequencing only tumour DNA achieved 100% sensitivity and 100% specificity relative to microsatellite FLA in 197 CRCs. smMIP-based sequencing of short MNRs fulfils other requirements of an ideal diagnostic test^26^, such as 100% classification concordance in repeat testing and robustness to common sample variables, including low MSI-high DNA content (LLoD approximated to 3%) and applicability to poor quality sample DNA from FFPE tissue. The assay LLoD was also superior to that of the MSI Analysis System (Promega). Furthermore, smMIPs incorporate molecular barcodes into reads allowing the number of sample DNA molecules sequenced to be quantified as a quality control metric. A minimum of 75 molecular barcodes per marker was required for accurate classification.

The inclusion of *BRAF* V600E in the smMIP-based MSI assay streamlines the LS screening pipeline, requiring only one tumour test prior to germline testing of MMR genes, equivalent to tumour-sequencing^22^. We were able to detect low variant allele frequencies in *BRAF* down to 1.7%, with improved sensitivity compared to HRM analysis, which has an estimated LLoD of 10%^33^. Using a ≥1% mutant read threshold, our frequency estimates of *BRAF* V600E in 9.4% and 36.6% of MMRp and MMRd CRCs agrees with the 7% and 31% frequencies previously observed^36^. *MLH1* promoter methylation is an alternative test for sporadic cases of MMRd CRC and has a higher specificity than *BRAF* V600E when screening for LS^37^. However, testing both markers is redundant due to their association^37^, and *MLH1* methylation testing reduces sensitivity for LS by deselection of *MLH1* mutation carriers that have methylation as a second hit^38^ or germline epimutations^39^. This was also observed by Hampel *et aß^2^*, and explains the reduced sensitivity of current screening practice relative to tumour-sequencing. We also multiplexed smMIPs targeting *KRAS* codons 12 and 13 mutations within our assay (Supplement S6). Whilst this does not cover the full scope of *RAS* gene mutations, it highlights the robustness of smMIPs to multiplexing and the ease with which additional, clinically actionable biomarkers can be added to the assay.

The cost of a diagnostic assay is a significant factor in its clinical uptake. Tumour-sequencing for example, has an estimated cost of 607±207€ per sample^24^, significantly more than a targeted assay such as our smMIP-based MSI assay. Our assay has an equivalent reagent cost to FLA when using 24 microsatellite plus *BRAF* markers, ranging from £8.20-£32.60 depending on the capacity of the MiSeq kit used (Supplement S7). As 6 microsatellites were sufficient for accurate MSI classification these costs can be reduced further. The smMIP protocol can also be fully automated and ported to the higher throughput NextSeq platform^40^ (Supplement S8). Furthermore, *BRAF* V600E testing is included within the assay, avoiding expenditure on additional tests for LS screening.

In summary, our smMIP-based MSI assay is highly sensitive and specific for MMR deficiency in CRC, simultaneously detects *BRAF* V600E, is reproducible, and is robust to sample variables. The automation of laboratory workflow and result interpretation removes the need for expert personnel, and provides a cheap, scalable assay. Combined, these factors suggest that a high throughput smMIP-based MSI assay is a suitable companion diagnostic for immune checkpoint blockade therapy and is applicable to two-step LS screening strategies.

## Acknowledgements

Thanks to Professor Majlinda Lako, Institute of Genetic Medicine, Newcastle University, UK, for gifting H9 DNA.

Thanks to Dr Marian Case and Professor Julie Irving, Northern Institute for Cancer Research, Newcastle University, UK, for gifting HCT116 and K562 cell lines.

Thanks to Dr Katharina Wimmer, Division of Human Genetics, Medical University of Innsbruck, Austria, for gifting control DNAs extracted from peripheral blood lymphocytes of consenting patients.

## Supplemental Data

Supplement S1: Contents of .bed file, listing marker loci used by MIPgen to generate single molecule molecular inversion probe sequences.

Supplement S2: Oligonucleotide sequences for single molecule molecular inversion probes targeting 24 microsatellite and *BRAF* V600E loci, amplification primer sequences, and custom sequencing primer sequences.

Supplement S3: Volumes of each single molecule molecular inversion probe used to “balance” (reduce variation in) the number of reads obtained from each marker, based on per marker read depths from the training cohort which used each probe at equal concentration.

Supplement S4: Formula to calculate the expected proportion of reads with a mutant microsatellite from sequencing mixtures of MSI-high DNA and MSS DNA, based on the proportions observed from sequencing pure MSI-high DNA and pure MSS DNA.

Supplement S5: Agarose gel of amplification products generated from variable input quantities of template DNA for 9 samples.

Supplement S6: Mutation frequencies from sequencing of *KRAS* codons 12 and 13, obtained by including an additional single molecule molecular inversion probe in the probe multiplex.

Supplement S7: Reagent costs per sample, depending on the capacity of the MiSeq kit used.

Supplement S8: Comparison of the proportion of reads with insertion-deletion mutations in all microsatellites from smMIP-based sequencing of 15 samples on the NextSeq platform versus the MiSeq platform.

